# An Isogenic Cell Line Panel for Sequence-based Screening of Targeted Anti-cancer Drugs

**DOI:** 10.1101/2021.08.06.455336

**Authors:** Ashley L. Cook, Nicolas Wyhs, Surojit Sur, Blair Ptak, Maria Popoli, Laura Dobbyn, Tasos Papadopoulos, Chetan Bettegowda, Nickolas Papadopoulos, Bert Vogelstein, Shibin Zhou, Kenneth W Kinzler

## Abstract

We describe the creation and characterization of an isogenic cell line panel representing common cancer pathways, with multiple features optimized for high-throughput screening. More than 1,800 cell lines from three normal human cells were generated using CRISPR-technologies. Surprisingly, we discovered most of these lines did not result in complete gene inactivation, despite integration of sgRNA at the desired genomic site. However, a subset of the lines harbored true, biallelic disruptions of the targeted tumor suppressor gene, yielding a final panel of 100 well-characterize lines covering 19 pathways frequently subject to loss of function in cancers. This panel included genetic markers optimized for sequence-based ratiometric assays for drug-based screening assays. To illustrate the potential utility of this panel, we developed a multiplexed high-throughput screen that identified Wee1 inhibitor MK-1775 as a selective growth inhibitor of cells with inactivation of *TP53*. These cell lines and screening approach should prove useful for researchers studying a variety of cellular and biochemical phenomena.

## Introduction

Recent advances in chemical synthesis techniques and robotics have led to an expansion in the availability of small molecule libraries [1, 2]. With the availability of curated libraries containing more than a million compounds, screening emphasis has shifted to identifying good targets and robust screens to efficiently exploit these libraries [3, 4]. High-throughput screening (HTS) assays can broadly be divided into biochemical and cell-based assays. Biochemical assays enjoy the advantages of low cost, facile scaling, specificity of measured outcome, and the ability to incorporate rigorous controls [5]. However, not all pathways, cellular functions or phenotypes can be adequately captured in biochemical assays. For example, cell-based assays have the advantage of directly identifying compounds that produce the desired biological effect via known or unknown mechanisms [6].

The unprecedented progress in defining the cancer genome gave rise to hope for the development of new targeted cancer therapeutics. This hope was largely driven by early success of targeted therapies that inhibited the function of oncogenic driver mutations [7]. However, while the typical adult solid tumor harbors 3 or more driver gene mutations, most of these mutations affect tumor suppressor genes, with many tumors lacking even a single oncogene mutation [8]. Even when effective therapies for targeting oncogenes are found, resistance to monotherapy is almost guaranteed in patients with major tumor burden [9–11]. The optimal strategy to overcome this resistance is to treat patients with combinations of drugs targeting different cancer growth mechanisms [9]. But as noted above, more than one oncogene mutation is unusual in most common cancer types.

Effective strategies for targeting loss of functions associated with tumor suppressor gene (TSG) mutations would substantially increase the number of therapeutically addressable pathways. Unfortunately, to date, only one FDA approved therapy specifically exploits a TSG loss of function mutation [12, 13]. This therapy, as well as other approaches for targeting the loss of function associated with TSG mutations is based on the concept of synthetic lethality or essentiality [14, 15]. This concept was originally described in yeast, and a key aspect of assigning specificity to synthetic lethality is the availability of isogenic cells differing only in a single genetic alteration [16, 17]. CRISPR-based technologies allow the creation of such lines in human cells.

In response to these issues, we created a human isogenic cell line panel targeting 19 critical genes inactivated in cancer. Each of these lines was engineered using CRISPR-based methods to disrupt a single tumor suppressor gene, and each contained a unique genetic barcode to permit multiplex screening. For each cell line, multiple orthogonal assays were used to validate successful gene disruption. Moreover, the panel was constructed from three distinct normal cell lines to ensure the generality of observed affects. And, finally, a sequence-based ratiometric assay was designed from this panel that incorporates numerous internal controls to maximize the reliability and sensitivity of the screening process.

## Results

### CRISPR-Cas9 Creation of the Isogenic Cell Line Panel Targeting Critical Tumor Suppressor Gene Pathways

We first sought to create a resource for screening compounds active in critical cancer pathways. We focused on 22 pathways that were collectively altered in greater than two-thirds of the cancers as assessed in multiple large scale sequencing efforts (Table 1 and S1). In total, 22 TSGs and 3 control genes with known small molecule sensitivities (Table S2) were chosen for targeting by CRISPR mediated knockouts (Table 1). Typically, 6 gRNAs were employed for each target gene (range 6 to 12) with three chosen from published studies and three designed de novo, targeting either known mutation sites identified from the COSMIC database or early exons within the gene [18, 19] (Table 1 and S3). A total of 162 gRNAs were individually introduced into lentivirus constructs for gene targeting. Three distinct non-cancerous epithelial cell lines, RPE1 (retinal), MCF10A (breast), and RPTec (renal), were targeted. All three lines have a predominately normal karyotype and the only known genetic alteration among the three lines was a homozygous deletion of the *CDKN2A* gene in MCF10A [20]. After transduction with lentiviral CRISPR-Cas9 and puromycin selection, over 1,800 individual CRISPR targeted single cells were picked and expanded for subsequent characterization.

**Table 1:**
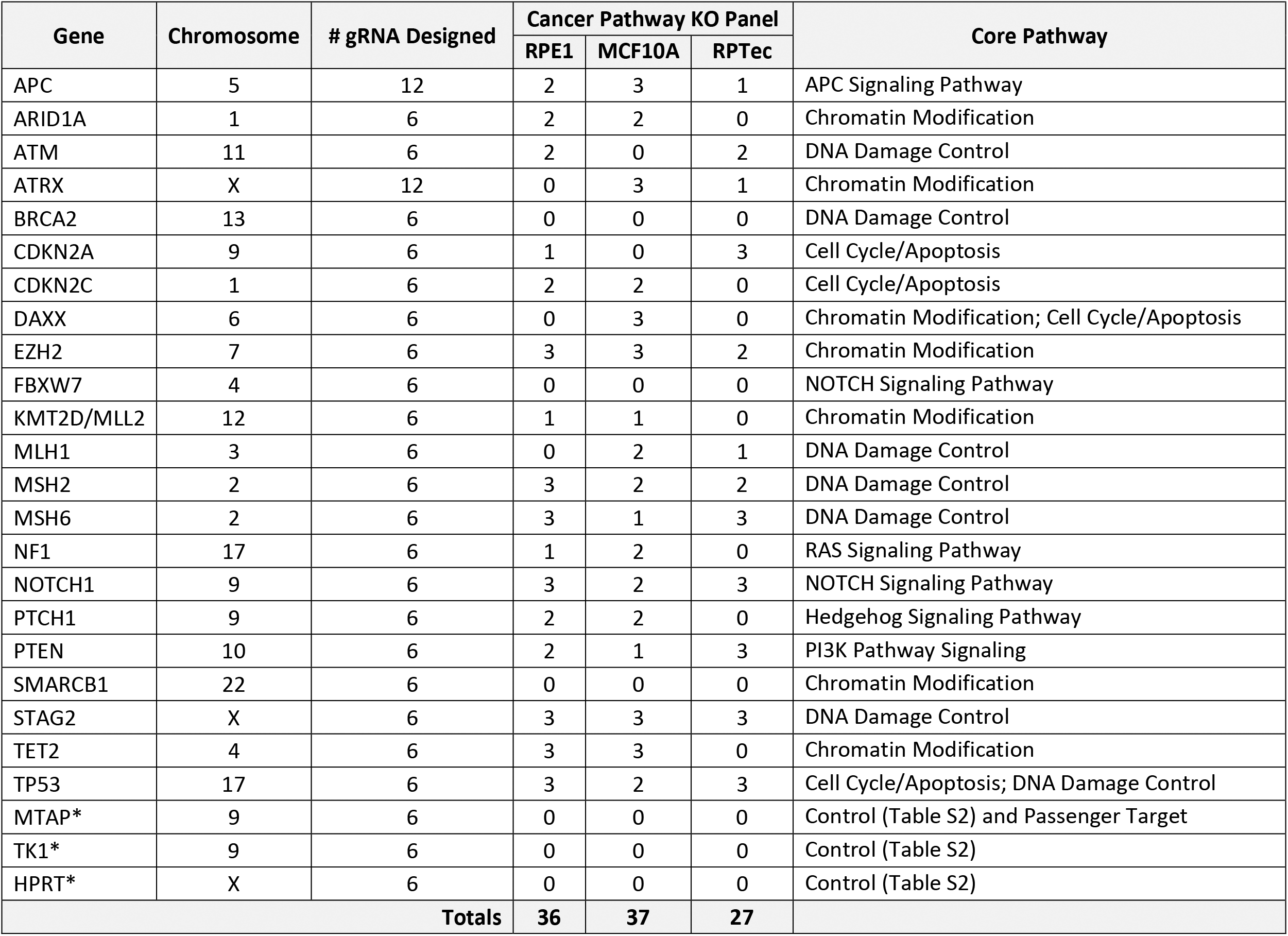
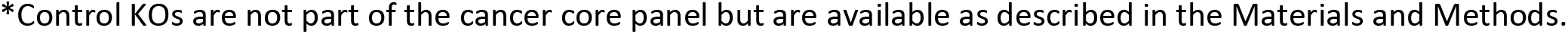
Cancer Pathway Knockout Panel. 100 cell lines composing the Cancer Pathway Knockout Panel detailed by targeted genes and cell line background. Details of gRNA can be found in Table S3. Cellular Processes with Core Pathways in parentheses were defined as in Vogelstein et al. [8].

| Gene | Chromosome | # gRNA Designed | Cancer Pathway KO Panel |  |  | Core Pathway |
| --- | --- | --- | --- | --- | --- | --- |
|  |  |  | RPE1 | MCF10A | RPTec |  |
| APC | 5 | 12 | 2 | 3 | 1 | APC Signaling Pathway |
| ARID1A | 1 | 6 | 2 | 2 | 0 | Chromatin Modification |
| ATM | 11 | 6 | 2 | 0 | 2 | DNA Damage Control |
| ATRX | X | 12 | 0 | 3 | 1 | Chromatin Modification |
| BRCA2 | 13 | 6 | 0 | 0 | 0 | DNA Damage Control |
| CDKN2A | 9 | 6 | 1 | 0 | 3 | Cell Cycle/Apoptosis |
| CDKN2C | 1 | 6 | 2 | 2 | 0 | Cell Cycle/Apoptosis |
| DAXX | 6 | 6 | 0 | 3 | 0 | Chromatin Modification; Cell Cycle/Apoptosis |
| EZH2 | 7 | 6 | 3 | 3 | 2 | Chromatin Modification |
| FBXW7 | 4 | 6 | 0 | 0 | 0 | NOTCH Signaling Pathway |
| KMT2D/MLL2 | 12 | 6 | 1 | 1 | 0 | Chromatin Modification |
| MLH1 | 3 | 6 | 0 | 2 | 1 | DNA Damage Control |
| MSH2 | 2 | 6 | 3 | 2 | 2 | DNA Damage Control |
| MSH6 | 2 | 6 | 3 | 1 | 3 | DNA Damage Control |
| NF1 | 17 | 6 | 1 | 2 | 0 | RAS Signaling Pathway |
| NOTCH1 | 9 | 6 | 3 | 2 | 3 | NOTCH Signaling Pathway |
| PTCH1 | 9 | 6 | 2 | 2 | 0 | Hedgehog Signaling Pathway |
| PTEN | 10 | 6 | 2 | 1 | 3 | PI3K Pathway Signaling |
| SMARCB1 | 22 | 6 | 0 | 0 | 0 | Chromatin Modification |
| STAG2 | X | 6 | 3 | 3 | 3 | DNA Damage Control |
| TET2 | 4 | 6 | 3 | 3 | 0 | Chromatin Modification |
| TP53 | 17 | 6 | 3 | 2 | 3 | Cell Cycle/Apoptosis; DNA Damage Control |
| MTAP* | 9 | 6 | 0 | 0 | 0 | Control (Table S2) and Passenger Target |
| TK1* | 9 | 6 | 0 | 0 | 0 | Control (Table S2) |
| HPRT* | X | 6 | 0 | 0 | 0 | Control (Table S2) |
| **Totals** |  |  | **36** | **37** | **27** |  |

\*Control KOs are not part of the cancer core panel but are available as described in the Materials and Methods.

### Genetic Characterization of Candidate Knockout Lines

A massively parallel sequencing approach was used to assess targeting and ensure that essentially all of the cells within any chosen cell line had the expected genotype. For this purpose, a SafeSeqS approach was implemented which utilizes unique molecular barcodes to reduce errors from PCR or sequencing [21]. For each gRNA, two distinct sets of primer pairs were designed to cover the targeted region. In total, 324 PCR primer pairs were designed and used to amplify the 162 gRNA genomic target regions (Table S4). This analysis confirmed successful gene disruption in only 302 of the greater than 1,800 lines tested. Though one might have expected a higher fraction of successfully targeted lines based on the previous successes of functional screens [22], our criteria for gene disruption were particularly stringent: both alleles had to contain out-of-frame insertions or deletions that could not be readily “rescued” by skipping an exon during splicing. Moreover, we used high depth sequencing and required that the fraction of reads containing an intact targeting site was < 1%. In the 302 lines chosen on the basis of the sequencing results, the deletions ranged from 1bp to 38bp and the insertions ranged from 1bp to 43 bp (Figure. S1). Over 31% of cell lines harbored a single base pair insertion or deletion, and an additional 9% of the lines harbored 2 bp insertion or deletion (Figure S1). The targeting success rate varied across genes and cellular backgrounds. Overall, we successfully identify cell lines with biallelelic gene inactivation in 22 of the 25 targeted genes in one or more cellular background, covering 50 of the theoretically possible 75 gene-cell line combinations (Figure 1A, Table S5)[23, 24].

**Figure 1.**
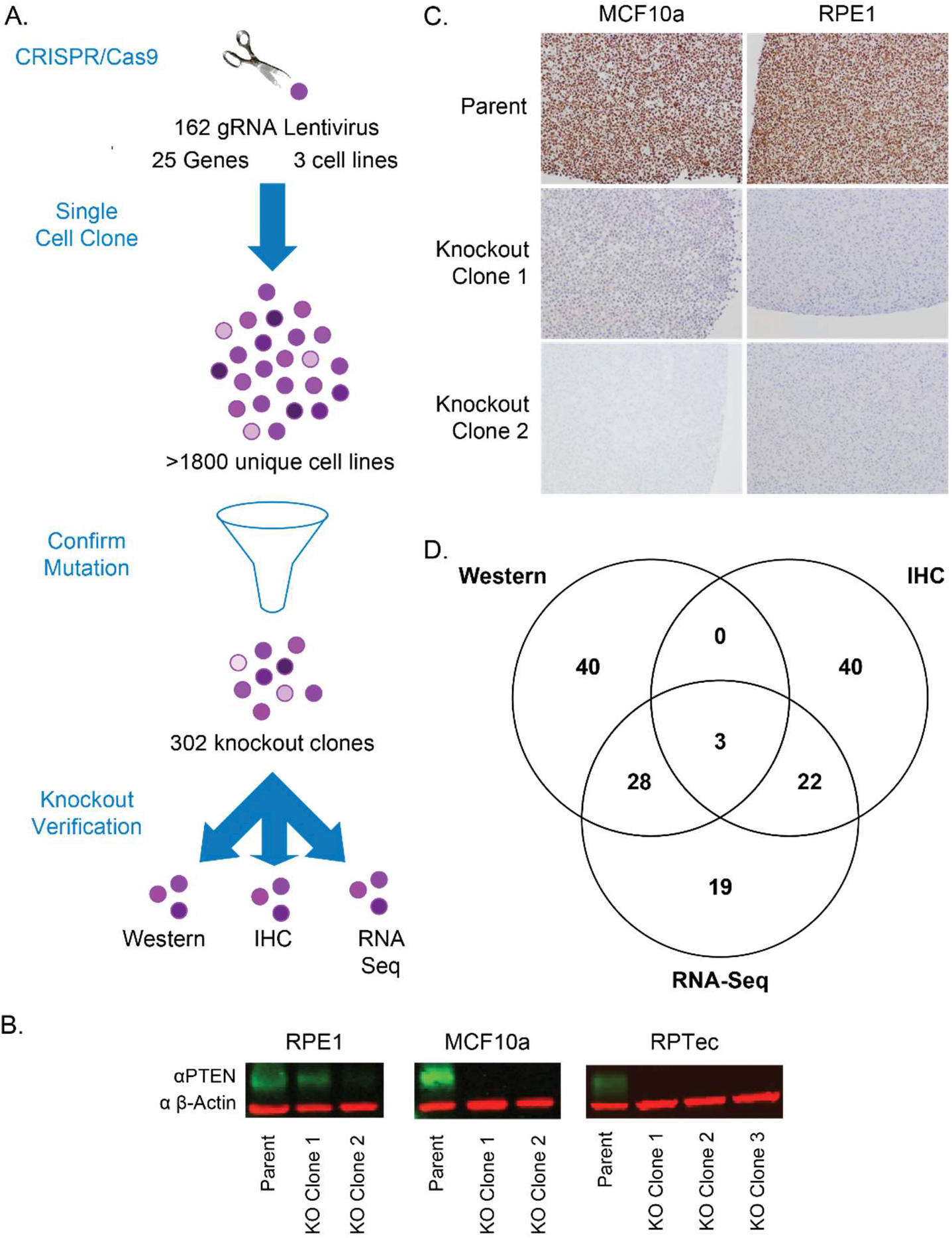
Design and Validation of Knockout Isogenic Cell Line Panel. A. Knockout cell lines were generated by targeting each gene individually with 6 or more gRNAs for a total of 162 gRNAs covering 25 genes in 3 cell lines (RPE1, MCF10a, and RPTec) (Table 1 and S3). Over 1,800 single cells were selected and expanded, then targeted NGS employed to verify bialleic out of frame insertion or deletion mutations, of which, 302 of these new-found cell lines met the criteria. B. Representative western blot expression of *PTEN* (green) and *?-actin* (red) in RPE1, MCF10a, and RPTec cell lines. The first knockout cell line in RPE1 is not a true knockout, with *PTEN* protein present, while the other knockout cell lines show no *PTEN* protein expression. C. Representative IHC staining of *ARID1A* protein in RPE1 and MCF10a cell lines demonstrating protein loss. D. 302 Cell lines with confirmed genetic mutation underwent secondary knockout verification of a combinatory of protein and/or RNA. A total of 152 cell lines passed protein and/or knockout validation with the number of cell lines passing each method indicated. Circles in Venn diagram are not drawn to scale to improve readability.

### Orthogonal Validation of Knockout Lines

We next sought to orthogonally validate the disruptions in these 302 lines. We established a hierarchical validation strategy where we first sought to establish loss of protein by western blot analysis, followed by immunohistochemistry (IHC) and finally loss of wild type transcript by transcriptome analysis.

Western blot assays were performed on 95 cell line and protein loss was confirmed in 71 of them (Figure 1B, Table S6). For 102 of the cell lines, we performed IHC assays and confirmed protein loss in 65 lines (Figure 1C, Table S6). Finally, to validate 4 genes lacking western or IHC assays and to begin to characterize the transcriptomes of additional selected lines, we constructed RNA-Seq libraries from 97 isogenic cell lines and sequenced them to an average depth of 2.2 x 10e7 reads per cell line (Table S6). To be validated by RNA-Seq, at least 15 reads (Average=87.5, N=19) covering the mutated or flanking exons were required with no evidence of wild type sequence or in-frame exon skipping. In total, we were able to validate loss of normal gene product at the protein or RNA level in 152 cell lines, representing 20 of the 22 targeted genes (Figure 1D, Table S5). One hundred of these 152 lines were subsequently assembled into the “Cancer Pathway Knockout Panel” to minimize overlap while maximizing diversity (Tables 1 and S7).

As noted above, several genes with known chemical sensitivities were also targeted to provide controls for assay development (Table S2, S8 and examples in Figure S2). In addition, we exploited the known differential sensitivity of cells without genetic inactivation of *TP53* to the small molecule MDM2 inhibitor Nutlin-3a[25]. Nutlin-3a causes cell senescence or death in cell lines with functional *TP53* by increasing the amount of available p53 protein. As expected, cell lines with wild type *TP53* were 5-10 times more sensitive to Nutlin-3a than their *TP53* null counterparts (Figure S3).

### Development of a multiplexed ratiometric cell growth assay

To demonstrate the potential utility of our engineered TSG knockout panel, we developed a screening platform that permits co-culture of multiple cell lines in a single well. Each of the cell lines in a single well thereby provides multiple internal controls for drugs that are generally toxic, rather than specifically toxic to a cell line harboring a specific disrupted pathway. Primers were designed to universally PCR-amplify every integrated gRNA in our cell line panel. Subsequent sequencing of the amplification products produced unique DNA barcodes for each cell line in a well. The representation of individual barcodes in the sequencing data thereby reflected the number of cells with the particular pathway disruption (Figure 2A). Though we constructed knockouts in three different parental cell lines, we pooled only cell lines derived from one parental cell line in any single well to more easily control for differences in parental cell line growth (Figure 2A).

**Figure 2.**
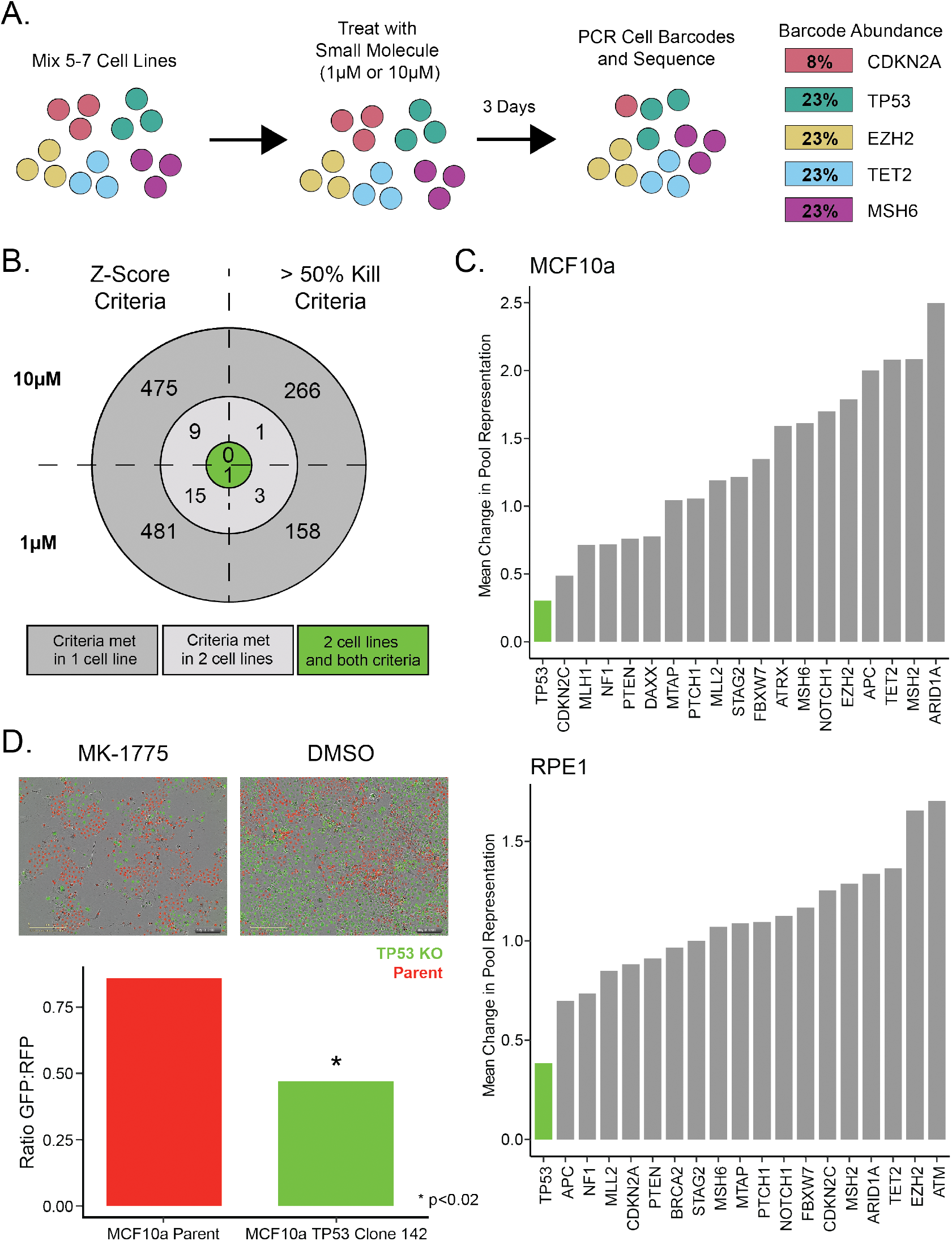
Synthetic Lethal Screen with FDA Approved Small Molecules. A. Cartoon example of high-throughput assay design showing hypothetical killing of a *CDKN2A* knockout cell line. B. Results summary for compounds of interest (gray) and hits (green) from the small molecule screen. C. Average change in fraction of reads for all knockout cell lines in RPE1 at 10μM and MCF10a at 1μM stratified by knockout gene in the presence of MK-1775. D. Co-Culture of TP53 wild type MCF10a parent (RFP) and *TP53* knockout cell line MCF10a TP53 142 (GFP) treated with MK-1775. A quantification of the image is shown as a bar graph.

With this pooling strategy (Fig 2A and S4), we could obtain nearly 15,000 measurements of cell growth from a single lane of an Illumina HiSeq 2500 instrument. This typically represented the output of 22 plates, each containing 96 wells using 5 to 7 cell lines per well. The multiple intra-well and inter-well measurements, along with positive and negative controls in every 96-well plate, provided an interlocking group of ratiometric measurements that enhanced specificity. Indeed, we observed a false positive rate of only 0.11% for the negative control well (DMSO only, no drug).

### High-Throughput Screen of FDA Approved and Clinical Trial Compounds

Using the multiplex assay described above, we evaluated a library of 2,658 FDA approved small molecule compounds for their ability to inhibit the growth of cell lines with specific pathway defects. For this screen, we used 81 cell lines derived from two different parental cell lines and representing 19 targeted pathways (See Table S5 for cell lines used in screen). Each of the 81 cell line were exposed to 1μM or 10μM of compound for 72 hours (Figure 2A). As negative controls, each 96-well plate included wells without drug or vehicle and wells with only vehicle (DMSO). The positive controls included in each 96-well plate were one well treated with Nutlin-3a (an inhibitor of normal p53 function) and one well treated with staurosporine (a non-specific, cytotoxic control). The negative and positive controls performed as expected, with DMSO having no effect on growth (Figure S5A) and staurosprine producing a marked reduction in growth (Figure S5B). The other positive control (Nutlin-3a) documented a pronounced difference between the growth of cell lines dependent on their *TP53* status, as expected (Figure S5C).

In total 430,596 compound-cell line interactions were scored in this assay. Compounds of interest were identified by requiring both a statistically (i.e., Z-score of −1.5 or less) and biologically significant (i.e., greater than 50% inhibition) effect as described in Material and Methods. Furthermore, it was required that this criteria were satisfied by independent cell lines with the same pathway disrupted in two parental cell backgrounds (Figure 2B). After applying these stringent filters to the 2,658 FDA approved small compounds, 1 hit emerged: *TP53* loss sensitized cells to the effects of MK-1775 (Figure 2B and 2C).

### TP53 deficiency sensitizes to MK-1775 (AZD1775)

MK-1775 demonstrated selective growth inhibition of *TP53* deficient cell lines from both RPE1 and MCF10a backgrounds within the primary screen when treated in the low micromolar ranges (Figure 2C). This results was confirmed in an orthogonal fashion using co-cultures of GFP (*TP53* mutant) and RFP (*TP53* wild type) labeled isogenic knockout cell lines (Figure 2D). MK-1775 is an inhibitor of *Wee1, a* kinase that controls the G2/M transition [26]. Previous studies have indicated that MK-1775 can selectively inhibit the growth of TP53 deficient cells in human cancers in vivo and in vitro in combination with radiation or chemotherapy and MK-1775 is currently in clinical trials for *TP53* deficient tumors in combination with chemotherapy or radiation [27–31]. Thus, our result does not represent a new drug discovery but rather represents an unbiased proof-of-principle for the new assay.

## Discussion

The results present above document two aspects of a novel resource for drug screening. First, we describe a panel of highly characterized isogenic cell lines containing single gene knockouts in critical cancer pathways. Second, we describe a multiplex, sequence-based assay that can be used for drug screening.

One important characteristic of our panel is the extensive validation undertaken for candidate knockout cell lines. Each of them had out-of-frame insertions or deletions which could not be “exon-skipped” without giving rise to a down-stream out-of-frame event. Moreover, all cell lines show a lack of functional RNA or protein products. In total, we derived a panel of 100 well-annotated isogenic cell lines that were validated in this way. Not all of the cell lines have to be included in a drug screen, particularly an initial one. But the redundancy inherent in the cell lines described here allows rapid confirmation of the activity of a drug identified in an initial screen. The variety of pathways and cellular backgrounds represented in these lines should provide an ideal resource for phenotypic high-throughput screening for a wide range of disease targets.

## Acknowledgements

The results shown in Table S1 are based upon data generated by the TCGA Research Network: https://www.cancer.gov/tcga. The authors would like to thank Dr. Sujayita Roy and Dr. Alan Meeker of the Johns Hopkins Sidney Kimmel Comprehensive Cancer Center Oncology Tissue Services Core for their assistance with IHC staining and optimization. Funding: This work was supported by the Virginia and D. K. Ludwig Fund for Cancer Research, the Lustgarten Foundation for Pancreatic Cancer Research, the Commonwealth Fund, the Bloomberg-Kimmel Institute for Cancer Immunotherapy, Bloomberg Philanthropies, the Mark Foundation for Cancer Research, and NIH Cancer Center support grant P30 CA006973.

## Author Contributions

NW, ALC, SS, KWK, NP, SZ, BV, and CB participated in the design and planning of the project. NW, ALC, BP, MP, and LD performed research NW, ALC, and KWK wrote the manuscript.

## Supplemental Materials

## Materials and Methods

### Cell Lines & Cell Culturing

RPE1, HEK293, MCF10a, and RPTec cells were purchased from The American Type Culture Collection (Virginia, USA). RPE1 cells were grown in RPMI 1640 Medium (Invitrogen, California, USA, Cat #11875-119) supplemented with 10% fetal bovine serum (HyClone, Utah, USA, Cat #16777-006). RPTec cells were grown in EPITHELIAL CELL MEDIUM-Complete Kit (Science Cell Research, California, USA, Cat #4101). HEK293 was grown in DMEM (Thermo Fisher, USA, Cat# 11995065) supplemented with 10% FBS (HyClone, Utah, USA, Cat #16777-006), MCF10a cells were grown in Bullet Kit MEBM Basal Medium 500 ml with MEGM SingleQuots Kit Suppl. & Growth Factors (Thermo Fisher, USA, Cat# CC3150). In vitro all cells were grown at 37°C with 5% CO_2_. Mycoplasma testing performed by The Genetic Resources Core Facility at Johns Hopkins University School of Medicine (Maryland, USA).

The cell line characterized in this study are detailed in Table S5. A subset of the lines were validated and banked for distribution including 100 lines targeting the 19 critical cancer pathways (Table S7) and 8 lines where control non-cancer pathway were targeted (Table S8). Of the non-tumor suppressor genes targeted, *MTAP* is of particular interest and represented by multiple lines because it is frequently co-deleted with *CDKN2A* making it passenger mutation targetable for therapeutic benefit in human cancers [32].

### CRISPR-Cas9

Integrated CRISPR-Cas9 gRNAs were designed using Chop-Chop based upon common mutations sites identified in COSMIC [33]. Each gene was targeted with 6-12 gRNAs (Table S1). gRNAs were ordered from IDT Technologies (Iowa, USA) with the addition of ligation sequences: caccgNNNNNNNNNNNNNNNNNNNN and aaacNNNNNNNNNNNNNNNNNNNNc. gRNAs were ligated into the LeniCRISPR V2 plasmid (Addgene, Massachusetts, USA, Cat #52961) using previously published protocol [18]. CRISPR/Cas9 plasmid was virally transduced into cells using Lenti-X Packaging Single Shots (VSV-G) using manufacturer’s instructions (Clontech, California, USA, Cat #631275). See Table S3 for list of all gRNAs utilized.

### Mutation Detection and Analysis

DNA was extracted from cells using Quick Extract (Lucigen, Wisconsin, USA, Cat #QE09050) and amplified using primer pairs listed in Table S4 designed to amplify 66-80 base pair segments containing the predicted cut site for each of our gRNAs listed in Table of gRNAs. Primer sets were designed for the SafeSeqS application, and were sequenced on an Illumina MiSeq and analyzed as previously described [34].

### Western Ab Information

Cells were lysed using RIPA buffer (Thermo Fisher, USA, Cat #89901) with x1 protease inhibitor (Thermo Fisher, USA, Cat #4693159001) and left on ice for 30min. Samples were then centrifuged at max speed for 3min in a QIAshredder (Qiagen, Maryland, USA, Cat #79654) before being transferred to a new Eppendorf tube. Protein was quantified using a BCA assay (Thermo Fisher, USA, Cat #23227).

Westerns were performed by loading 30-50ug of total protein per well into 15 well polyacrylamide gels (Bio-Rad, California, USA, Cat #456-1086) and run for 30min at 200V. Gels were then transferred using manufacturer’s instructions (based on size) to nitrocellulose membrane using a Bio-Rad turbo transfer apparatus. Membranes were blocked for 1hr with 3% milk TBS-Tween before being incubated overnight in primary antibody (concentration dependent on antibody). Membranes were then washed 4 times for 5min each with TBS-Tween. Secondary antibody was applied at 1:2500 (Abcam, United Kingdom, anti-mouse Cat #ab6728 or anti-rabbit Cat #ab6721). Membranes were imaged using Pierce™ ECL Western Blotting Substrate (Thermo Fisher, USA, Cat #32106) following manufacturer’s instructions and imaged on a Bio-Rad Chemidoc (Bio-Rad, California, USA). Table S9 indicated antibodies used for westerns and their typically employed concentrations.

### Immunohistochemistry (IHC)

Antibody concentrations for IHC were specific for each protein being screened (Table S9). Generally, Immunolabeling for a protein was performed on formalin-fixed, paraffin embedded sections on a Ventana Discovery Ultra autostainer (Roche Diagnostics, Switzerland). Briefly, following dewaxing and rehydration on board, epitope retrieval was performed using Ventana Ultra CC1 buffer (Roche Diagnostics, Switzerland, Cat #6414575001) at 96°C for 64 minutes. Primary antibody was applied at 36°C for 60 minutes. Primary antibodies were detected using an anti-rabbit HQ detection system (Roche Diagnostics, Switzerland, Cat #7017936001, #7017812001) followed by Chromomap DAB IHC detection kit (Roche Diagnostics, Switzerland, Cat #5266645001), counterstaining with Mayer’s hematoxylin, dehydration and mounting.

### RNA Seq Methods

For RNA extraction, cells were pelleted, frozen in liquid nitrogen, and stored at −80°C until RNA extraction. RNA extraction using Qiagen AllPrep DNA/RNA Mini Kit (Qiagen, Maryland, USA, Cat# 80204) per manufacturer’s instruction with cell homogenization in RLT buffer via QIAshredder (Qiagen, Maryland, USA, Cat# 79656). RNA quality control using Agilent Tapestation 2200 (Agilent, California, USA, Cat# G2964AA) and the Agilent RNA ScreenTape (Agilent, California, USA, Cat# 5067-5576) and Agilent RNA ScreenTape Sample Buffer and Ladder (Agilent, California, USA, Cat# 5067-5577, Cat# 5067-5578) per manufacturer’s instruction. Library prep using Illumina RNA library prep kit (Illumina, California, USA, Cat # RS-122-2001) and sequenced on an Illumina HiSeq 4000 paired end using manufacturer’s instructions.

Sequencing reads aligned to Hg38 using HISAT2 (version 2.0.5), assembly and quantification was performed using StringTie (version 1.3.3) and differential expression was performed using R package Ballgown (version 2.6.0) as described [35]. Exon skipping was determined using IGV Viewer Sashimi Plots [36].

### Statistical Testing

Z-scores calculated utilizing the ratio of target cell UID to total sequencing reads in a well

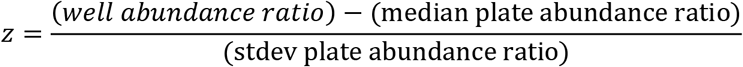

### Small Molecule Cell Line Screen

We identified optimal plating of cells up to 7 distinct cell lines from the same background and 5,000 total cells per well of a 96-well plate maintained the best cell line representation and compound response. The library chosen was the FDA-approved & Passed Phase I Drug Library in 96 well format (Selleck Chem LLC, Houston TX, #L3800). We screened 81 total knockout cell lines across MCF10a and RPE1 cell line backgrounds and 19 critical cancer pathways at two doses: 1μM and 10μM. Cells were plated in the morning and treated with the compound libraries in the evening of day 1. Plates were harvested on day 4 and molecular barcodes identifying each cell line were quantified by high-throughput sequencing.

Sequencing using barcoded forward primer including Illumina primer sequence, N14, plate barcode, and LentiV2 sequence ( AATGATACGGCGACCACCGAGATCTACACTCTTTCCCTACACGACGCTCTTCCGATCT NNNNNNNNNNNNNN**BARCODES**TGTGGAAAGGACGAAACACC). Reverse primer includes Illumina primer sequence, well barcode, spacer, and a LentiV2 sequence (CAAGCAGAAGACGGCATACGAGAT**BARCODES**NNCGGACTAGCCTTATTTTAACTTGC). This amplicon requires 96 reverse primers and 1 forward to amplify and uniquely identify each well in a 96-well plate (Figure S4). In total 25 plate and 192 well barcodes were designed and verified. These primers were used to amplify cell pools after DNA extraction using Quick Extract (Lucigen, Wisconsin, USA, Cat #QE09050). Amplified reads were sequenced on either an Illumina MiSeq or Illumina HiSeq 2500.

Screen controls were scored by evaluating the ratio of unique identifier (UID) reads matching a single cell line in each staurosporin treated well to the cell lines respective UID reads from the untreated DMSO wells in the same screen plate. This was performed for each knockout line within each screen plate. To assess cell line representation and performance in each treated well, we calculated the z-score for each cell line’s fraction of reads within a single well to compare cell line abundance in drug treated wells and compared to the 95 other wells within the same plate. We assume the null hypothesis – for any given compound, it will not have a specific interaction with our gene of interest. Compounds in our screen with UIDs less than two-fold more unique molecular barcodes than the non-specific small molecule control within the same plate, staurosporine, were classified as non-specific cell killing. We did not consider compounds from these wells in our analysis.

To determine the z-score threshold we looked at representation of each cell line in the DMSO treated control wells and down-sampled the sequencing of these well to 0 in increments of 10%, calculating the z-score at each increment to determine the power to see each cell line. This was determined for each plate in our screen and thresholds determined from the 3^rd^ quartile, the lowest 25% of plates based on cell line representation within the plate. Based on this in silico calculated 3^rd^ quartile z-score we classified cell lines as well powered, and used a cutoff of −1.5, or low powered, and used a cutoff of −1.0 to identify compounds of interest. Based on this classification, a maximum z-score threshold was set for each cell line either −1.5 for well powered or −1 for low powered. All compound-cell line z-scores below these thresholds were considered compounds of interest. For a compound to be considered a hit, the majority of cell lines in two cell line backgrounds would need to identify it as a compound of interest. Applying these criteria to DMSO wells showed that 0.11% of controls wells met the hit criteria.

### Hit Validation – Fluorescent Labeled Cell Lines & Cell Confluence

*TP53* knockout cell lines were labeled with either a GFP or RFP plasmid (Essen Bioscience, USA, Cat #4477 and #4478). We then performed a co-culture of each knockout cell line with its respective parental line and treated with the compound of interest in a dose response curve. We imaged cells every 6 hours for 4-6 days of treatment using Incucyte Zoom (Sartorius, Michigan, USA). Fluorescence was quantified using four locations in each treated sample by the IncuCyte Zoom 2016B software.

Confluence assays were plated in 96-well plate format and imaged every 6 hours for 4-6 days of treatment using Incucyte Zoom.

### Hit Validation – Sybr Green Cell Counting Assay

Cell response assays were quantified using a sybr readout. Cells were rinsed 2x in phosphate buffered saline (PBS) (Thermo Fisher, USA, Cat#10010-049) and lysed with 50μL of 0.2% SDS (Thermo Fisher, USA, Cat#15553027) for 2 hours at 37°C. 150μL of Sybr Green I (Thermo Fisher, USA, Cat#S7563) solution (1:750 in water) was added and mixed 10x with a pipette. Fluorescence was read out with 485nm excitation and emission measured at 530nm on a BioTek microplate reader (BioTek, Vermont, USA). DNA content of each sample analyzed relative to untreated samples.

---

**Supplemental Figure S1:**
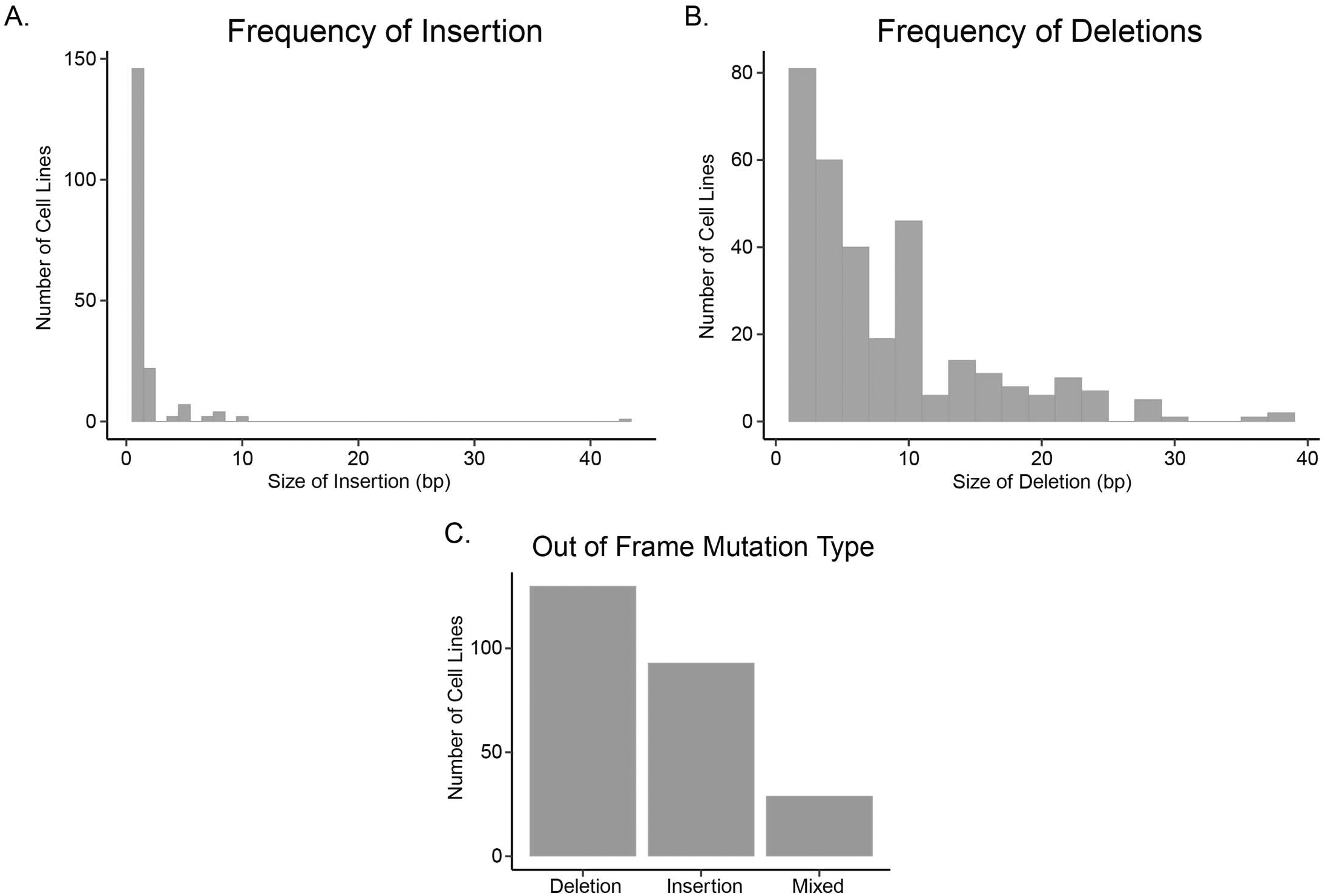
Histogram of insertions A. and deletions B. present in cell lines with biallelic out of frame mutations identified by SafeSeqS. The x-axis shows the number of bases inserted or deleted in bins of 2bp. C. Number of cell lines containing specific mutation types. Mixed denotes cell lines with an out of frame deletion in one allele, and out of frame insertion in the other.

**Supplemental Figure S2:**
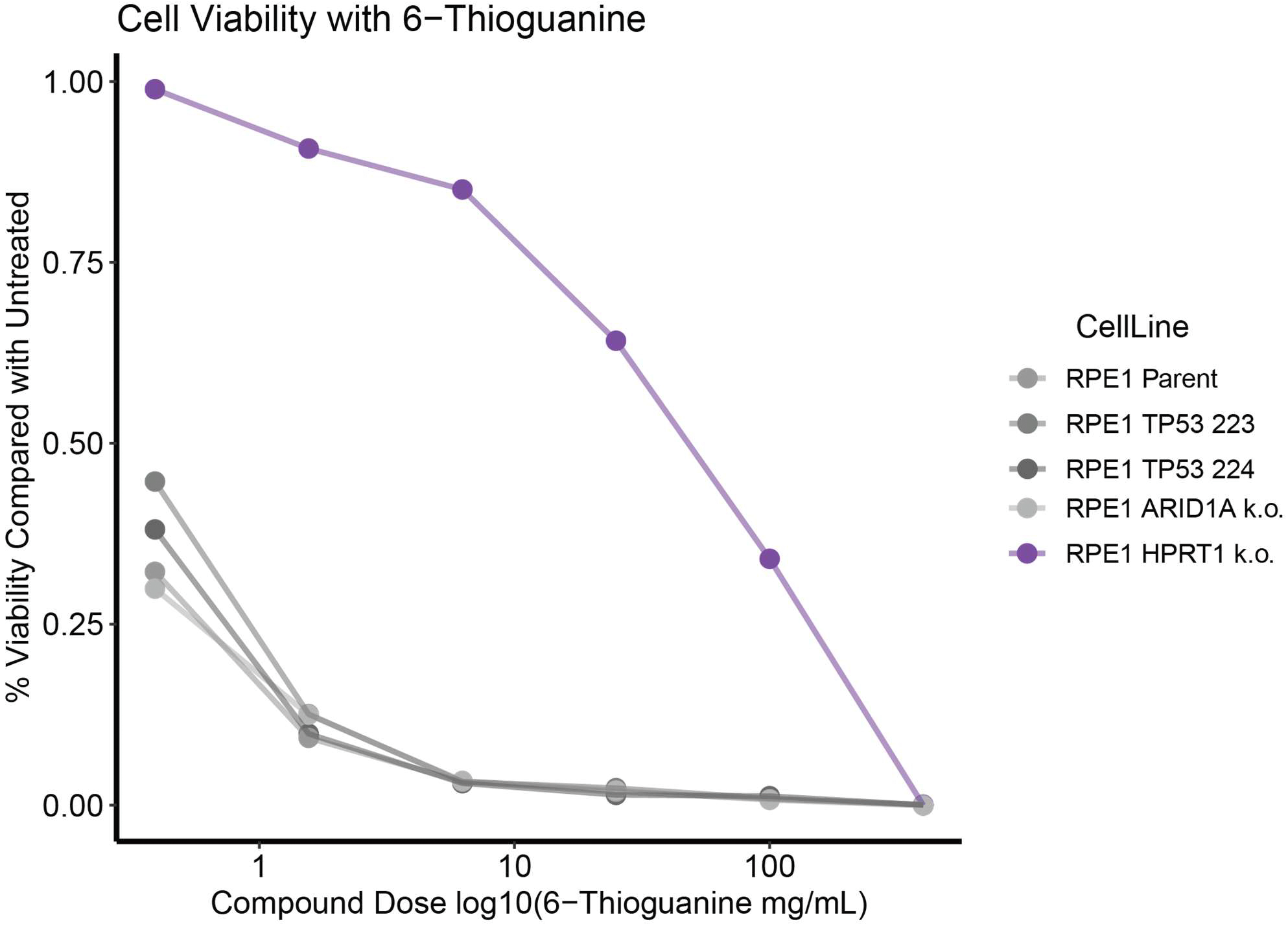
*HPRT1* and *TP53* Knockouts from the RPE1 background treated with 6-Thioguanine (6-TG) for 3 days and readout by SYBR green assay (DNA content).

**Supplemental Figure S3:**
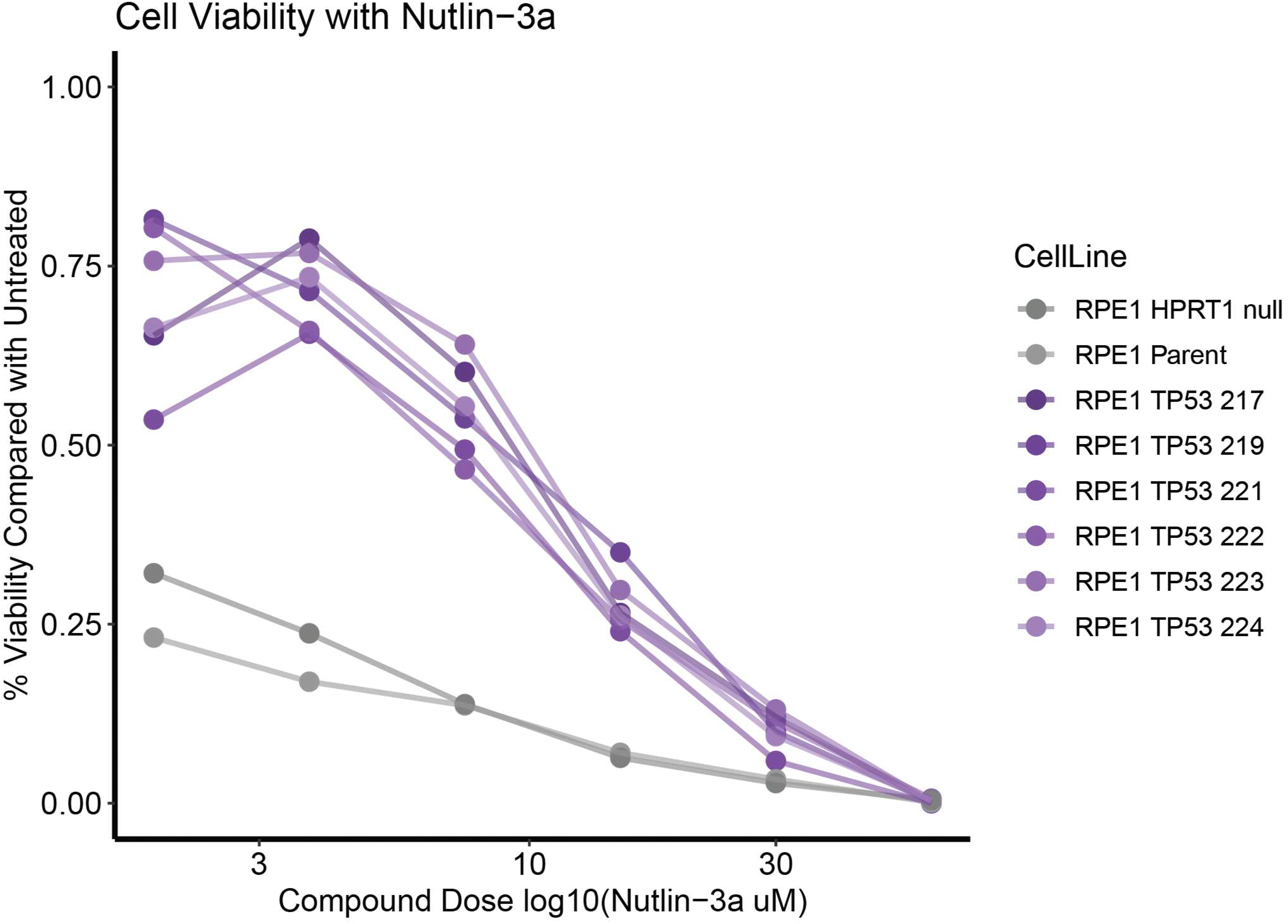
Dose response of RPE1 *TP53* knockout cell lines compared with RPE1 parent and RPE1 *HPRT1* knockout cell lines to Nutlin-3a, treated over 3 days and readout by SYBR green assay (DNA content).

**Supplemental Figure S4:**
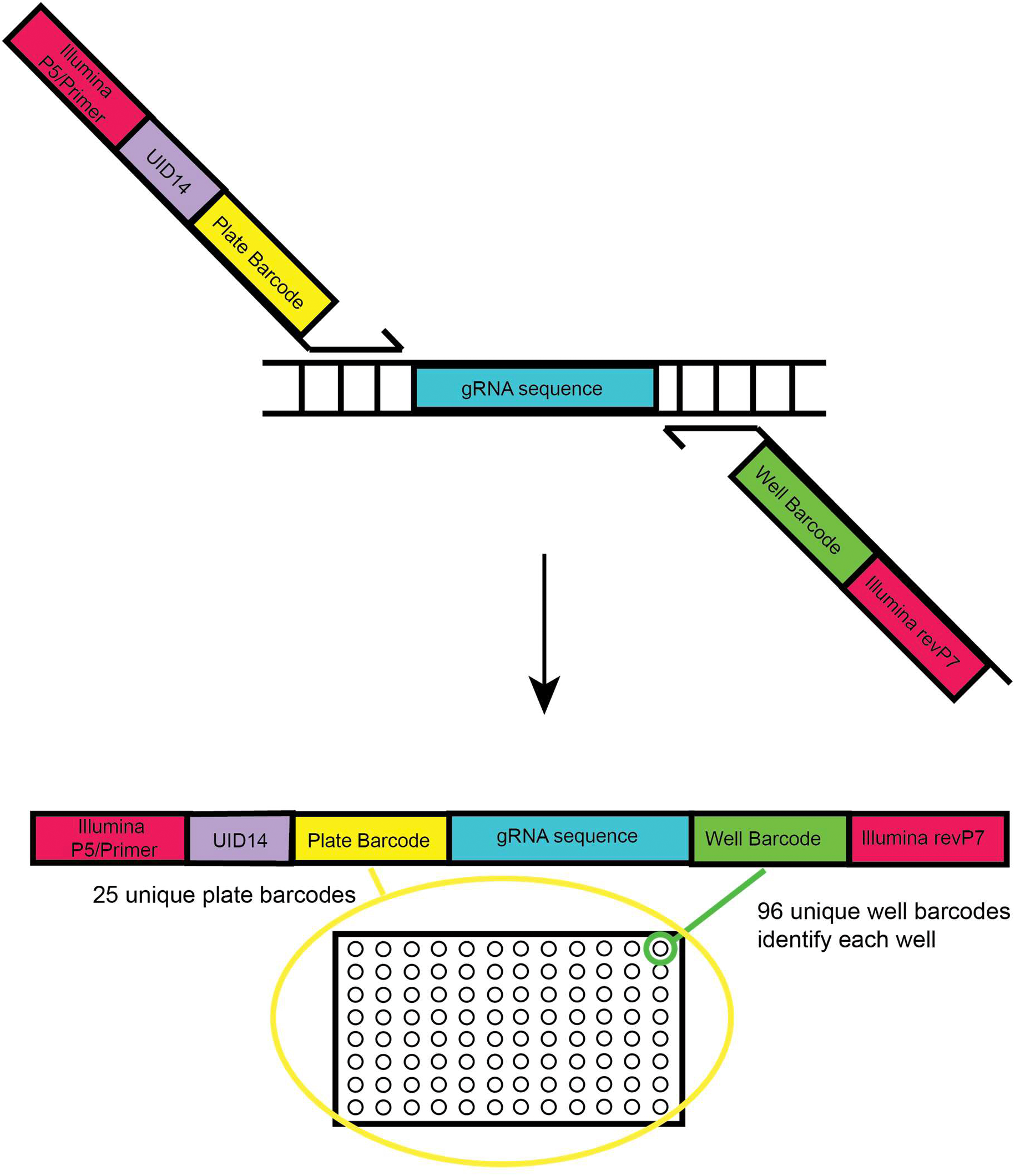
Cartoon showing amplification strategy of integrated barcode (gRNA) for high throughput sequencing. Up to 7 clones were per well were co-amplified with the above strategy at a time (each clone contains a unique barcode gRNA). The amplicon is 197bp in size.

**Supplemental Figure S5:**
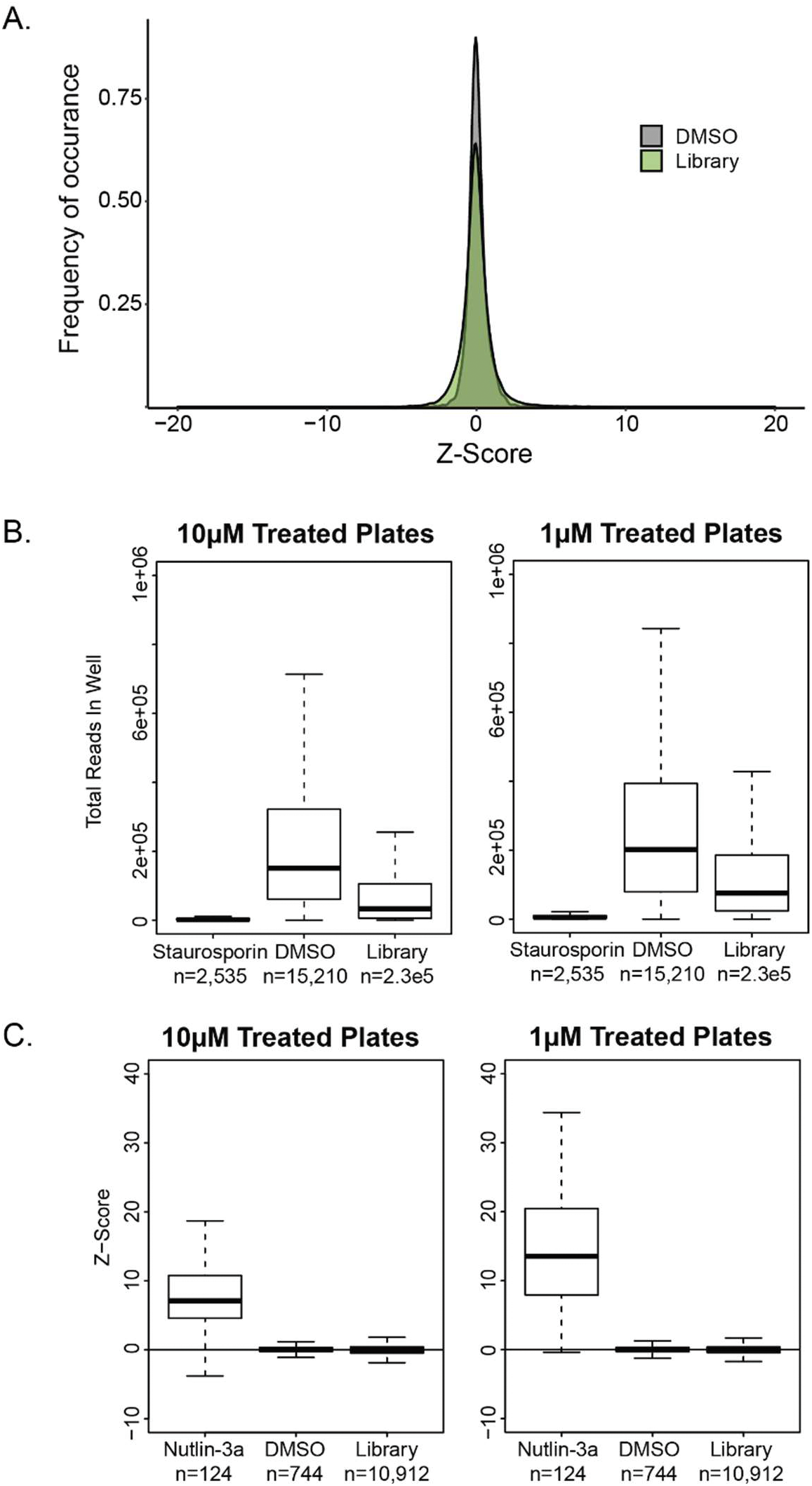
A. Distribution of Z-Factors in the Selleck Chem HTS for DMSO control wells and wells treated with a compound (library). B. Boxplot showing total number of unique identifier reads in control or library containing wells. C. Boxplot showing distribution of Z-Factors for *TP53* knockout clones when treated with DMSO, library or nutlin-3a. Z-Factors were calculated based on plate based median and standard deviation, excluding control wells.

